# Multiscale Analysis of Cellular Senescence through Ripley’s Functions and Functional Statistics

**DOI:** 10.64898/2026.06.29.734207

**Authors:** Cécile Verrier, Vanessa Dehennaut, Sophie Dabo-Niang

**Affiliations:** Univ. Lille, CNRS, INRIA, UMR 8524 – Laboratoire Paul Painlevé, F-59000 Lille, France; Univ. Lille, CNRS, Inserm, Institut Pasteur de Lille, CHU Lille, UMR9020-U1366 – CRCLille – Cancer Research Center of Lille, F-59000 Lille, France

**Keywords:** Cellular senescence, Functional Data Analysis, Ripley’s L function, Modeling, Spatial analysis

## Abstract

Cellular senescence is a heterogeneous and evolving process involved in development, tissue repair, aging, and age-related diseases. Although senescence burden in tissues has been widely studied, its spatial organization remains poorly understood, particularly in vivo. Senescence encompasses a spectrum of distinct states, with cells differing in molecular signatures, secretory activity, persistence, and interactions with their microenvironment depending on the inducing stimulus and tissue context. This heterogeneity suggests that spatial organization may reflect underlying processes such as tissue repair, regeneration, or maladaptive remodeling, providing insight into senescence function and its pathological roles. Here, we propose a quantitative, multi-scale framework to characterize the spatial organization of senescent cell populations in post-infarction mouse hearts. By combining a senescence-signature scoring strategy with spatial statistical methods and functional data analysis, we assess whether senescent cells exhibit clustered or dispersed patterns, and how these spatial distributions evolve over time following infarction. This approach aims to provide new insights into the spatiotemporal dynamics of senescence in vivo and to identify spatial features that may inform therapeutic strategies targeting age-related and tissue repair-associated pathologies.

## Introduction

Cellular senescence is a fundamental biological process involved in development, tissue repair, aging, and age-related diseases including cardiovascular diseases, neurodegenerative disorders and cancers [1, 2]. It is characterized by a prolonged cell cycle arrest accompanied by profound phenotypic changes, including metabolic rewiring and the acquisition of a senescence-associated secretory phenotype (SASP) [3, 4, 5], which can exert both beneficial and detrimental effects depending on the context [4]. Senescence can be triggered by a wide range of cellular stresses, including telomere attrition, genotoxic stress, oncogene activation, oxidative stress, mitochondrial dysfunction, and therapeutic insults such as chemotherapy or radiotherapy [3, 6]. This diversity of inducing stimuli contributes to the heterogeneity of senescent cell states across tissues and disease settings. Rather than representing a uniform endpoint, senescence encompasses a spectrum of distinct cellular states that differ in molecular signatures, secretory activity, persistence, and interactions with the microenvironment. These states may display functional plasticity depending on the inducing stimulus and tissue context, suggesting that senescence should be viewed as a dynamic and context-dependent process rather than a fixed phenotype [3].

For example, emerging evidence indicates that metabolic regulation can influence therapy-induced cell fate decisions in proliferating cells. In particular, modulation of specific metabolic pathways can prevent the establishment of a senescence program and instead promote apoptotic cell death [7]. While senescence plays beneficial roles in development, wound healing, and tumor suppression, its chronic persistence and accumulation contribute to tissue dysfunction and the progression of age-related diseases. Consequently, senescent cells are increasingly considered as therapeutic targets, and strategies aimed at their elimination or modulation have emerged as promising approaches in several pathological contexts [8, 9, 10].

Despite their biological and clinical relevance, the in vivo spatial organization of senescent cells within tissues remains poorly understood. It’s unclear whether senescent cells are spatially clustered, randomly distributed, or organized in structured microenvironments that evolve over time. Addressing this question is essential, as spatial organization may reflect mechanisms of local propagation, microenvironmental regulation, or coordinated tissue remodeling.

Recent advances in spatial transcriptomics now allow senescence-associated signatures to be mapped in situ [6]. In this context, methods based on global spatial autocorrelation, such as Moran’s index, have been used to assess whether senescent signals exhibit non-random spatial distributions. For instance, previous work applied Moran’s index to spatial transcriptomics data from post-infarction mouse hearts, revealing evidence of spatial clustering of senescence-associated signals at certain time points [6]. However, such global statistics provide only a limited description of spatial organization, as they do not capture the spatial scale of interactions and may mask more complex, multi-scale structures.

To overcome these limitations, spatial point process methods offer a powerful framework for analyzing the spatial arrangement of cells at multiple scales. Among these, Ripley’s K-function [11] and related summary statistics are widely used to characterize spatial patterns such as clustering or inhibition across distances [12, 13]. These approaches have been extensively applied in oncology and spatial biology [14], to study tumor architecture, immune infiltration, and cell–cell interactions. However, their application to the study of cellular senescence remains limited, particularly in the context of high-resolution spatial transcriptomics data.

Moreover, existing analyses of Ripley’s functions are often restricted to pointwise or descriptive interpretations, that do not fully exploit their functional nature. Functional Data Analysis (FDA) [15] provides a natural framework to adress this limitation by treating spatial summary functions as functional observations. This enables the characterization of entire spatial interaction profiles and the identification of underlying structures through techniques such as functional principal component analysis and clustering [16, 17].

Here, we propose a statistical framework that combines spatial point process methodology and Functional Data Analysis to investigate the spatial organization of senescent cells. Using spatial transcriptomics data from post-infarction mouse hearts and senescence signature scoring, we model the distribution of senescent cells as spatial point patterns and characterize their organization using Ripley’s functions and their variants. These functions are then analyzed within a functional data framework to identify dominant modes of spatial variation and to classify spatial patterns across conditions and time points.

To the best of our knowledge, this is the first study to integrate multi-scale spatial statistics and Functional Data Analysis for the characterization of cellular senescence in situ. This approach provides a quantitative framework to investigate the spatiotemporal organization of senescent cells and may help clarify their role in tissue remodeling and disease progression.

## Methods

### 2D Mouse Heart cell Data

The analysis presented in this work relies on a multi-step pipeline combining senescence scoring, spatial statistics and functional data analysis. Starting from Visium spatial transcriptomic data, hub-specific senescence scores are first computed and then used to characterize the spatial organization of senescent spots across tissue sections. The main steps of this workflow are summarized in Fig 1.

**Figure 1.**
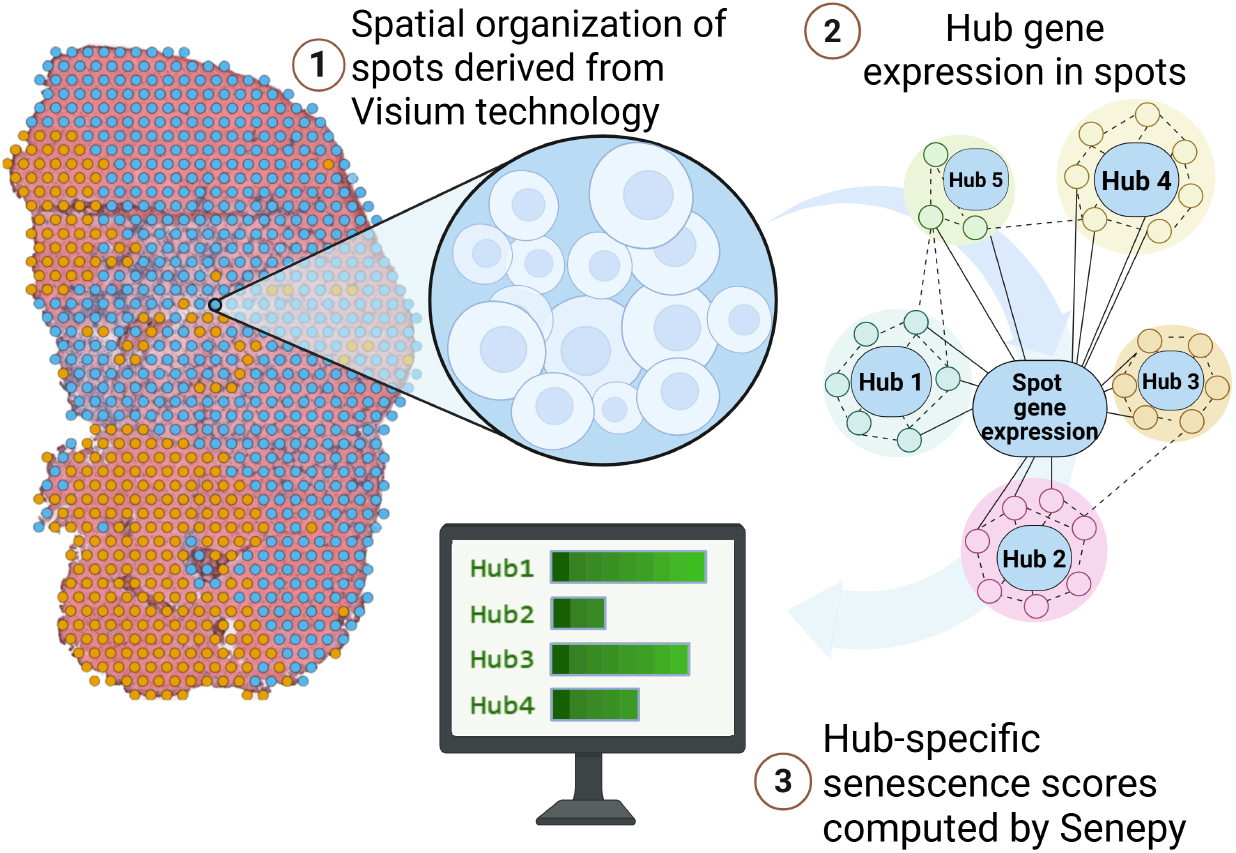
Workflow for the computation of hub-specific senescence scores. Schematic overview of the computation of hub-specific senescence scores from Visium spots using SenePy. Created in BioRender. verrier, C. (2026) https://BioRender.com/nmh8eal

The spatially resolved transcriptomic data used here come from the mouse heart dataset GSE176092 [18], obtained from the technology 10x Genomics Visium and made publicly available as a resource for the research community. This dataset contains heart sections collected at different times after infarction. All analyses were carried out directly on the spots of these Visium images, including the use of the biological hubs defined in the SenePy framework [6]. Unlike the study by Sanborn et al.[6] which integrates multiple single-cell datasets from different tissues, our work relies exclusively on this spatial dataset of mouse hearts to examine the spatial distribution of senescent spots in an infarction model.

The spatially resolved transcriptomic data from this dataset are provided as preprocessed and aligned AnnData objects, including gene expression counts per spot as well as the Visium spatial coordinates. These coordinates are given in pixels that we can convert to micrometers using the Visium specifications (spot diameter 55 *µm*), which in this case gives a conversion factor of *α* ≃ 2, 34 *µm/*pixel. The biological identities of the tissue regions (hubs) are directly available in the original publication. The raw data were not reprocessed (no fastq alignment or cell QC), because the Visium spots had already been filtered and aggregated by the authors of the dataset. All subsequent processing was therefore carried out directly from the normalized expression matrices in Scanpy and the associated annotations. During the preparation of this manuscript, Large Language Models (LLMs) were used to assist with language refinement and editorial improvements. All generated content was reviewed, verified, and approved by the authors.

To estimate the presence of senescent cells in tissues, we used SenePy, an open-source tool developed by Sanborn et al. [6] from transcriptomic signatures derived from single-cell data. Their approach relies on an unsupervised method that integrates prior biological knowledge according to which senescent cells accumulate with age but remain a minority in tissues. By analyzing the transcriptomic profiles of 43 murine cell types covering a wide age range, the authors identified dynamic genes specific to each cell type. From these genes, co-expression networks were constructed to reveal hubs, representing gene modules potentially associated with senescence, where each gene can be associated with a weight reflecting its importance within the network (e.g. through a number of edges). Genes not connected to these hubs were considered as noise or stochastic age-related signals, not directly relevant to cellular senescence. This approach led to the definition of 72 murine signatures associated with a given cell type. Although these signatures are highly heterogeneous, an enrichment in classical senescence markers was observed across several cell types.

In our pipeline, SenePy is used as a signature-scoring tool that assigns to each observation (here, each Visium spot) a quantitative score per hub, in the spirit of module-scoring approaches such as those described by Wolf et al. [19]). For each hub, SenePy provides both the signatures (gene lists, optionally weighted) and associated metadata. An identifier-matching step is then applied when necessary to harmonize the gene symbols present in the AnnData object with those expected by the hubs. The score of a hub is then computed, yielding for each spot a quantitative value representing the activity of the signature. Scoring is performed at the spot level rather than at the single-cell level, with the goal of obtaining a continuous measure of the relative expression of senescence hubs on the Visium grid of each section. Let ℋ_*h*_ ⊂ ℝ^2^ be a compact hub domain *h* = 1, …, *H*, equipped with Borel *σ*-algebra *ℬ*(ℋ_*h*_). For each spot *i* in ℋ_*h*_, let **x**_*i*_ ∈ ℋ_*h*_ ⊂ ℝ^2^ be the precise 2D coordinates, discrete mRNA counts 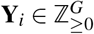 for *G* = 387 genes, and an assigned cell-type label *τ*_*i*_ ∈ *{*1,…, *K}*. The senescence (SenePy) score is computed as:

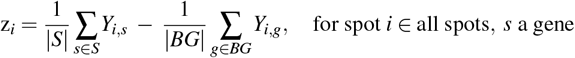

where |*S*| and |*BG*| denote the cardinality (number of elements) of the gene sets *S* and *BG*, respectively, where *S* corresponds to the set of genes belonging to the selected SenePy senescence signature (or hub), and *B*_*k*_ corresponds to a group of genes with similar mean expression levels, constructed in order to select comparison genes with expression levels close to those of the genes in the signature *S* (with the aim of preventing the score from being biased by global differences in gene expression levels). Here *Y*_*i,s*_ and *Y*_*i,g*_ represent the (optionally amplified) expression values of genes in the gene signature *S* and in the background gene set *BG*, respectively. A detailed description of the scoring procedure (including gene binning, background selection, and optional binarization and weighting) can be found in Sanborn et al. [6].

The general principle of the SenePy score is to compare the average expression of the hub genes with that of a background gene set selected in a controlled way based on expression levels, in order to limit biases related to the distribution of mean expression values. Options also make it possible to binarize the data and to amplify expression by assigning importance weights to genes, reflecting their level of connectivity within the co-expression network defining the hub. In our case, the continuous scores produced by SenePy are used as an intermediate step to define a senescence variable that can be exploited in spatial analysis, as described in the following subsection.

Consider in the following that the coordinates, count expressions, cell types and senescence scores are observations of the marked spatial point process {(**x**_*i*_, **Y**_*i*_, *τ*_*i*_, *z*_*i*_)} defined on probability space (Ω, ℱ, ℙ), where Ω denotes the set of possible outcomes, ℱ a *σ*-algebra of measurable events, and ℙ the associated probability measure. The process is defined over the spatial domain ℋ_*h*_, with locations **x**_*i*_ ∈ ℋ_*h*_ ⊂ ℝ^2^ and mark space 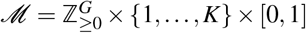, where the three mark components correspond to **Y**_*i*_, *τ*_*i*_, *z*_*i*_ ∈ [0, 1].

### Spatial scenesence point pattern

#### Moran’s index

In order to summarize the global spatial organization of a signal measured over a set of spots, it is common to use a spatial autocorrelation statistic. Moran’s index, introduced in 1950 [20], is one of the most commonly used measures for this purpose. We introduce an indicator function allowing the selection of pairs of points separated by a distance close to *r*:

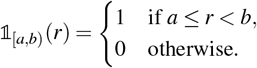

Using this indicator function, we formulate the Moran’s index used in this paper as:

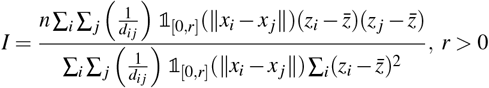

with *n* denoting the total number of spots, *x*_*i*_ = (*u*_*i*_, *v*_*i*_) and *x* _*j*_ = (*u*_*j*_, *v* _*j*_) represent the spatial coordinates of spots *i* and *j*, respectively, where the distance between two spots is given by the Euclidean norm 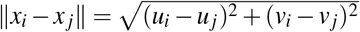. The indicator function **1**_[0,*r*]_(∥*x*_*i*_ − *x* _*j*_∥) is equal to 1 if the Euclidean distance between *x*_*i*_ and *x* _*j*_ is less than or equal to *r* (in our setting *r* is taken as 0), and 0 otherwise. Finally, *z*_*i*_ and *z* _*j*_ denote the senescence scores at spots *i* and *j*, respectively, while 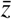 represents the empirical mean of the observed values.

Although widely used, Moran’s index remains a global measure that does not account for the variability of spatial organization according to the scale considered, hence the importance of complementing this global approach with finer measures of spatial organization.

#### Ripley functions

Proposed in 1976 [11], Ripley’s function is an indicator widely used to characterize correlations in point processes. In the context of our study, it allows the analysis of the spatial structure of senescent cells identified through their transcriptomic *SenePy* score. The principle of the *K* function relies on the calculation, for a given *r*, of the average number of events (here, senescent spots) present within a disk of radius *r* centered on a given point, normalized by the global density of the process. This density, called the *intensity*, is denoted *λ*, and corresponds to the average density of events in the study region. The analytical definition of the function is the following:

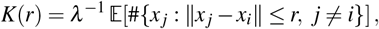

*λ* denotes the intensity of the point process and the expectation (E(.)) concerns the cardinality of the set [*x* _*j*_ : ≤ |*x* _*j*_ − *x*_*i*_| *r, j* ≠ *i*[that is, the number of points of events at a distance less than or equal to *r* from a typical point of the process. In practice, this function is estimated empirically from a finite sample of points observed in a spatial window *W*. The standard Ripley estimator is then written:

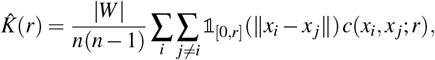

where |*W*| the area of the spatial window and spatial coordinates and Euclidean distances are as previously defined. The indicator function 1_[0,*r*]_(∥*x*_*i*_ − *x* _*j*_ ∥) is equal to 1 if the Euclidean distance between *x*_*i*_ and *x* _*j*_ is less than or equal to *r*, and 0 otherwise. Finally, *c*(*x*_*i*_, *x* _*j*_; *r*) denotes an edge correction factor intended to compensate for the underestimation of the number of neighbors near the boundaries of *W*. More precisely, as described by Marcon et al [21], Ripley’s correction assigns a larger weight to neighbors located close to the boundary according to the proportion of the neighborhood that remains observable within *W*.

The *K* function is said to be cumulative, because it integrates all distances less than or equal to *r*. The estimation of the function is carried out incrementally :

1. For each point, and for each distance *r*, the number of neighbors located in a circle of radius *r* is counted;
2. For each distance *r*, the average number of neighbors over all points is then computed, weighting each pair (*x*_*i*_, *x* _*j*_) by the edge correction factor *c*(*x*_*i*_, *x* _*j*_; *r*);
3. These results are compared with those expected under a homogeneous spatial Poisson model, which models a completely random distribution of events, in order to detect possible deviations (aggregation or dispersion).

This approach therefore makes it possible not only to detect a non-random organization of senescent cells, but above all to identify the relevant spatial scales at which this organization manifests.

#### Stationarity

Under the assumption of stationarity and a homogeneous Poisson process (that is, a spatial process without interaction between events), the theoretical reference function becomes:

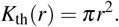

In other words, in a uniform space without particular spatial structure, the growth in the number of neighbors with distance simply follows the area of a disk. To facilitate graphical interpretation and obtain a linear representation under the null hypothesis of a homogeneous Poisson process, the Ripley transformation called the **Ripley** *L* **function** is often used:

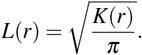

In the ideal case of a homogeneous CSR, we then have *L*(*r*) = *r*, which leads to representing *L*(*r*) − *r* in order to center the curve around zero in the absence of particular spatial structure; the sign of this deviation is then interpreted; *L*(*r*) − *r >* 0 indicating an excess of neighbors at distance *r* (aggregation) whereas *L*(*r*) − *r <* 0 indicates a relative deficit of neighbors (dispersion), and *L*(*r*) − *r* ≃ 0 an organization compatible with the null hypothesis, keeping in mind that in non-homogeneous or constrained configurations (irregular window, missing regions, discrete support), the reference curve may deviate from the theoretical line *L*(*r*) = *r*. Unlike Moran’s index which provides a single value, Ripley’s function offers a **multi-scale analysis**: it indicates not only whether senescent cells are clustered, but also from which distance these clusters are observed. This makes it possible, for example, to identify typical aggregation radii (tissue influence zones, specific microenvironments) or to compare spatial profiles between different biological conditions or cell types. This approach is particularly relevant because senescence is a heterogeneous phenomenon. The analysis using Ripley’s function therefore constitutes an appropriate tool to explore local dynamics, and to relate them to the functional organization of the studied tissue.

#### Pair correlation function (PCF) and variants

Unlike Ripley’s function, which accumulates interactions up to a distance *r*, the pair correlation function (PCF), denoted *g*(*r*), allows the evaluation of spatial interactions at a specific distance. In the following, we adopt the notation conventions introduced by Bull et al. [22]. In our case, 1_[0,*dr*)_(|*x*_*i*_ − *x* _*j*_ |− *r*) = 1 if the distance between points *i* and *j* belongs to the interval [*r, r* + *dr*), and 0 otherwise. The pair correlation function (PCF) is defined as:

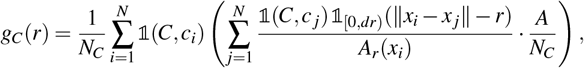

where *g*_*C*_(*r*) measures the average density of cells of type *C* located at a distance *r* from a spot of the same type, where the distance between two spots is defined as previously by the Euclidean norm.

*N*_*C*_ then represents the number of spots of this type and 1(*C, c*_*i*_) = 1 indicates whether spot *i* is of type *C*, 0 otherwise, where *c*_*i*_ denotes the category associated with spot *i*. Thus, for each spot *i* of type *C* in the observation domain Ω of area *A*, we define the annulus centered at *x*_*i*_ as the region located at a distance between *r* and *r* + *dr* from cell *i*, the division by *A*_*r*_(*x*_*i*_) then allowing the normalization of the number of points by the area of the annulus lying inside the domain, in order to obtain a comparable density even for points close to the boundary.

Thus, *g*_*C*_(*r*) *>* 1 indicates aggregation of spots of type *C* at distance *r, g*_*C*_(*r*) *<* 1 indicates dispersion, and *g*_*C*_(*r*) = 1 indicates a spatially random distribution.

In addition to the PCF, it is possible to localize spatially an interaction defined at a given distance *r* in order to identify the regions of the tissue that contribute the most to it. The topographical correlation map (TCM) thus makes it possible to spatially localize the regions of the tissue where the interaction is stronger or weaker than that expected under the hypothesis of complete spatial randomness.

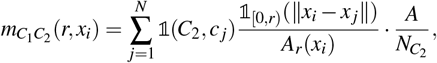

such that 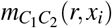 counts, for each spot *i* of type *C*_1_, the number of spots of type *C*_2_ located at a distance at most *r*. The division by *A*_*r*_(*x*_*i*_) plays the same role as for *g*_*C*_(*r*): normalizing the number of spots by the surface area effectively contained within Ω. The multiplication by 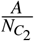 is then performed in order to compare this density to the average density of spots of type *C*_2_ in the domain.

The interpretation therefore remains the same as for the PCF: 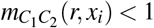 indicates dispersion and 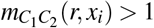 indicates aggregation between spots of type *C*_1_ and *C*_2_ at a distance at most *r*. To facilitate interpretation, 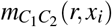 is normalized into a bounded variable 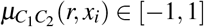 such that *µ* = 0 corresponds to the behavior expected under CSR, and such that correlation and exclusion are comparable on a linear scale.

The final map is obtained by spatial smoothing of the contributions 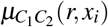 using a Gaussian kernel of width *σ* :

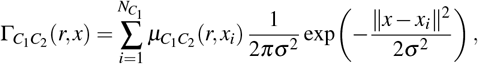

where the same notation conventions as previously introduced are used.

Positive values of Γ indicate regions where the interaction is stronger than expected under CSR, whereas negative values indicate local exclusion. The parameter *σ* controls the trade-off between spatial resolution and smoothing.

#### Spatial organization of senescent spots

For each Visium section, senescent spots are considered as a set of points in the plane, allowing their spatial distribution to be modeled as a point process observed on a finite tissue domain. The objective is to quantify the spatial aggregation of senescent spots as a function of distance. A Ripley curve is estimated for each sample, and then compared according to the time post-infarction. The calculation relies on the spatial coordinates of the spot centers provided by the Visium platform, and distances are evaluated on a predefined grid of radii. The observation window corresponds to the region effectively sampled after spot filtering, which may cause an irregular geometry and introduce edge effects.

#### Construction of a null distribution by permutations (conditional CSR)

On a Visium slide, spatial positions do not correspond to a homogeneous continuous plane but to a discrete set of sites (spot grid) restricted by a tissue mask (spots outside the tissue removed) and sometimes by internal “holes” related to section quality or filtering, so that, even in the absence of biological aggregation, the theoretical reference of a homogeneous point process under *Complete Spatial Randomness* (CSR) is not necessarily the line *L*(*r*) = *r*, and a deviation from this line may simply reflect the effective geometry of the observed domain and the grid constraint rather than a real interaction. In this context, a robust way to define a CSR reference is to construct an empirical null hypothesis by permutation of labels [13], while preserving the admissible positions (all valid spots) as well as the total number of senescent spots, and then randomly permuting the binary label “senescent / non-senescent” on these positions and recalculating *L*(*r*) at each iteration. Repeated *B* times, this procedure provides an empirical distribution:

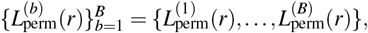

where 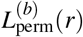 is the value of *L*(*r*) computed at distance *r* after the *b*-th permutation of the labels, and the set 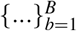 gathers the *B* realizations obtained. Thus, an average curve over the permutations can be obtained via :

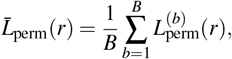

with, if necessary, an envelope obtained by quantiles.

The associated permutation *p*-value at distance *r* is defined by

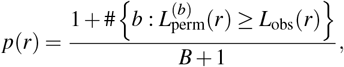

where *L*_obs_(*r*) denotes the observed value of the Ripley statistic at distance *r*.

#### Bootstrap of Ripley curves

As explained previously, in our study each biological sample (tissue section) provides a spatial curve *L*_*i*_(*r*) describing the organization of senescent spots as a function of the distance *r*. The entire dataset can thus be represented by a collection of functions:

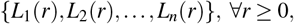

where *n* denotes the number of available samples. In our case, this number remains relatively limited, which makes the estimation of the statistical variability of the mean curves or derived statistics delicate. In order to evaluate the uncertainty associated with the estimates obtained from this reduced set of samples, we use a **non-parametric bootstrap** procedure. This approach consists in artificially generating replicates of the sample by performing resampling with replacement among the observed curves. More precisely, for each bootstrap replication *b* = 1, …, *B*, a new artificial dataset of size *n* is constructed by sampling with replacement from the set of observed curves:

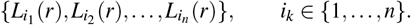

For each bootstrap dataset, the statistics of interest (for example the mean curve, the functional median, or certain characteristics derived from the curves) are recalculated. Repeating this procedure *B* times thus provides an empirical distribution of the estimates, making it possible to evaluate their variability and, if necessary, to construct confidence intervals. This approach does not modify the available biological information, but makes it possible to estimate more robustly the uncertainty associated with conclusions drawn from a limited number of independent samples.

#### Functional representation of the curves

For each sample *i*, the Ripley curve *L*_*i*_(*r*) is evaluated on a discrete grid of *m* = 100 equally spaced distances over [0, *r*_max_], with *r*_max_ = 250 pixels (≈ 607 µm). This bound remains smaller than the size of the tissue sections observed across the different samples, ensuring the reliability of the edge-corrected estimator. In order to manipulate them within a functional framework and to make the curves comparable across samples, they are represented as smoothed functions in a B-spline basis defined on a common distance domain. Each curve is thus approximated by a linear combination of basis functions:

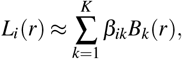

where 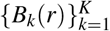 denotes a B-spline basis and *β*_*ik*_ the coefficients associated with sample *i*. This representation makes it possible to obtain curves defined on a common domain while reducing the noise related to the empirical estimation of the Ripley functions.

The spatial curves obtained for each sample can then be considered as observations of a functional process defined on a continuous domain corresponding here to the interval of spatial distances. In this framework, functional data analysis (FDA) consists in treating each observation as a function rather than as a discrete vector of values.

Formally, each Ripley curve *L*_*i*_(.), evaluated on a finite number of discrete grid points, is modeled as a realization of a functional random variable defined on the same probability space (Ω, ℱ, ℙ) introduced previously. An observation then corresponds to a function :

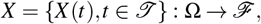

where *T* denotes a continuous domain and ℱ a space of functions, typically a Hilbert space such as *L*^2^(*T*). For each realization *ω* ∈ Ω, one obtains a function *t* ↦ *X* (*ω, t*), corresponding to an observed curve. In this framework, one can define statistical quantities adapted to functional data, in particular the functional mean:

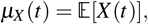

as well as the covariance function:

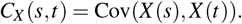

In practice, these quantities are estimated from a finite set of observed curves {*X*_1_, …, *X*_*n*_} using their empirical estimators, in particular the empirical functional mean:

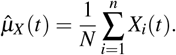

In our application, the observations *X*_*i*_ correspond to the smoothed Ripley curves *L*_*i*_(*r*) computed for each biological sample.

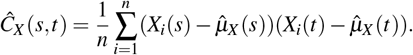

This framework makes it possible to describe the mean structure of the curves as well as their variability. In particular, the covariance structure can be analyzed using the spectral decomposition of the covariance operator, which constitutes the foundation of functional principal component analysis, as developed and applied in the context of spatial transcriptomic data by Dabo-Niang and Frévent [17]. More precisely, one considers the eigenfunctions *f* _*j*_(*t*) associated with the eigenvalues *λ*_*j*_ of the covariance operator *C*_*X*_, defined by the equation:

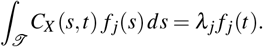

The eigenfunctions *f* _*j*_(*t*) describe the main directions of variation of the curves, and each observation can then be represented by its functional scores in this basis. This spectral decomposition constitutes the foundation of functional principal component analysis (functional PCA), which makes it possible to summarize the variability of the curves using a reduced number of eigenfunctions and their associated scores.

## Results

### Multi-scale spatial analysis of senescence

#### Detection of aggregation regimes using Ripley’s function L(r)

Senescence scores were computed from publicly available spatial transcriptomic data of post-infarction mouse hearts (DOI: 10.1038/s44161-022-00140-7) using the SenePy framework [6] (see Methods for details). These data consist of Visium spatial transcriptomic sections collected at multiple time points after myocardial infarction. The complete set of resulting spatial senescence score maps is available in Supplementary Information; two representative examples are shown in Fig 2 (a) and (b). These maps constitute the input for downstream analyses, including visualization of senescence patterns and multi-scale spatial characterization.

**Figure 2.**
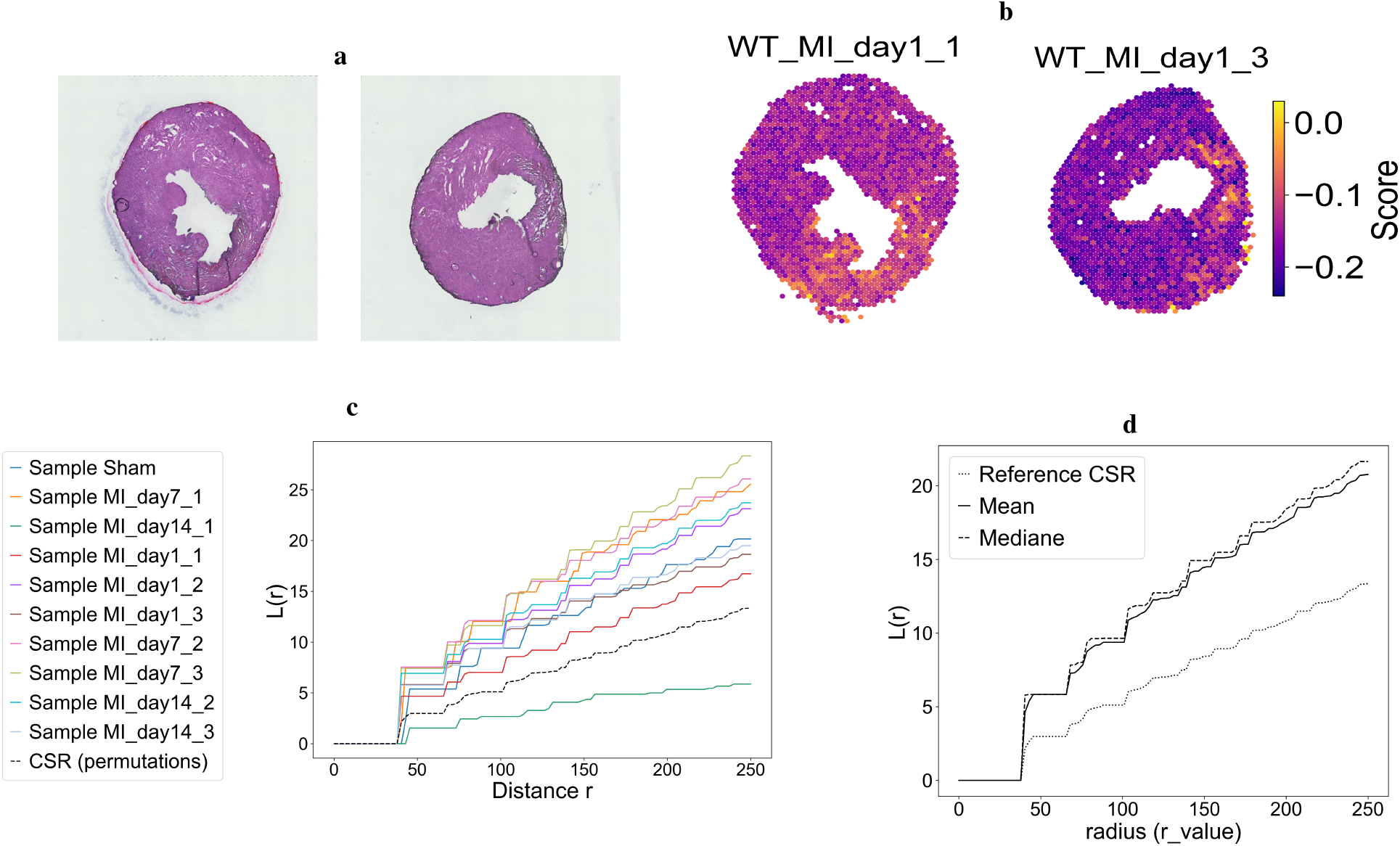
Exploration of the spatial structuring of senescent cells. **(a)** Representative heart sections from two biological replicates at one day post-infarction. **(b)** Spatial distribution of senescence scores identified using SenePy in the same samples; color intensity reflects hub-specific senescence scores across Visium spots. Moran’s index is reported to quantify global spatial autocorrelation. **(c)** Ripley’s L(r) functions computed for theses samples over 100 spatial scales (0 ≤ *r* ≤ 250 pixels). Each curve describes the spatial organization of senescence-associated spots as a function of distance r. The empirical CSR reference was obtained using 199 permutations of senescent/non-senescent labels (see Methods for details). **(d)** Mean and median Ripley’s L(r) curves aggregated across the samples, together with the corresponding mean CSR reference curve derived from the same permutation framework

The joint analysis of the mean and median curves of the *L*(*r*) statistic in Fig 2 highlights a spatial profile that is globally similar across samples. Indeed, the curves representing the mean and median values of *L*(*r*) in Fig 2 (d) remain well above the reference line. This suggests the presence of strong spatial aggregation on average from one sample to another, indicating that senescent spots exhibit a non-random organization and tend to cluster into aggregates rather than being diffusely distributed across the tissue section.

A change in trend can also be observed around a radius of 40 pixels, corresponding to approximately 94 *µ*m, where an inflection of the curves toward higher values becomes visible. This transition arises from the geometric constraint of the Visium slide: spots are arranged on a discrete grid and the center-to-center spacing between neighbors is typically on the order of 38–43 pixels, so that below this threshold there are no spot centers within the neighborhood and the statistic cannot explicitly count pairs of points. Thus, the portion of the curve for *r* below this minimal spacing mainly reflects the discretization of the support rather than a biological structuring, and the interpretation should therefore focus on distances beyond this threshold. Similarly, when examining the dynamics of *L*(*r*) separately for each sample in Fig 2(c), this aggregation between senescent spots persists.

Indeed, the Ripley curves lie largely above the reference curve for almost all samples, including both post-infarction samples and the sham sample, with p-values that are mostly significant (see Supplementary Tables S1 and S2). The only exception is the sample corresponding to a mouse heart section collected 14 days after myocardial infarction (replicat 2), which does not show clear spatial aggregation among senescent spots. This indicates an aggregated spatial organization of senescent spots relative to the hypothesis of complete spatial randomness for most of the distances considered. This spatial structure is already detectable in sham samples, suggesting that the observed organization is not exclusively induced by myocardial infarction but may reflect an intrinsic spatial patterning of senescence-associated transcriptional programs in cardiac tissue. Consistent with these results, Moran’s (I) values reported in Supplementary Table S2 indicate a significant positive spatial autocorrelation for the majority of samples, whereas one sample (second replicate at day 14 post-myocardial infarction) exhibits a Moran’s (I) value close to zero together with a non-significant p-value, suggesting a spatial distribution compatible with complete spatial randomness. Overall, these results indicate that senescence-associated transcriptional programs are spatially structured in cardiac tissue and that this organization is largely preserved across experimental conditions, with limited temporal variation and one divergent late-stage sample.

#### Bootstrap validation of the mean and median curves

Although the analysis of our curves already allows the identification of potentially atypical spatial profiles, a bootstrap based on resampling of the *L*(*r*) curves was performed in order to quantify the robustness of the aggregated profiles and to ensure that the resulting aggregated profiles are not dominated by a restricted subset of samples.

The bootstrap analysis presented in Fig 3 provides additional information on the variability observed between samples. The relatively wide uncertainty bands around the mean curve indicate that the intensity of senescent spot aggregation varies substantially between samples. This variability is consistent with the differences observed between the individual L(r) curves in Fig 2. Nevertheless, despite this heterogeneity, the overall spatial profile remains remarkably stable: the curves generally remain above the CSR reference and exhibit similar aggregation regimes across spatial scales. These results suggest that the previously identified aggregation pattern is robust to resampling and is not driven by a small subset of samples.

**Figure 3.**
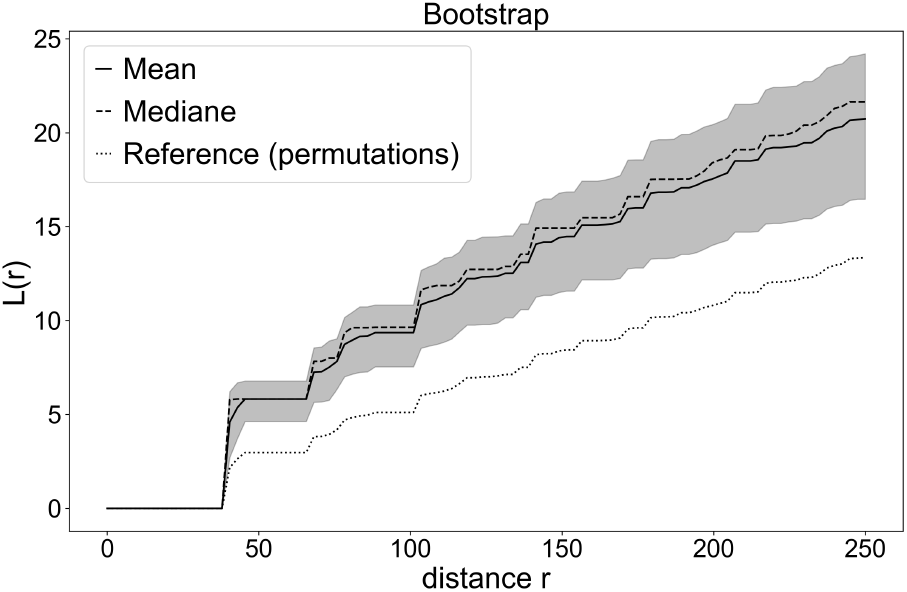
Bootstrap resampling of the mean and median of the Ripley *L*(*r*) statistic. The solid curve and the dashed curve correspond respectively to the mean and median of *L*(*r*) estimated by bootstrap (*B* = 1000 bootstrap resamples obtained by sampling curves with replacement). The shaded region indicates the 95% bootstrap confidence interval, computed under the same settings as in Fig 2.

### Functional analysis of Ripley curves

#### Smoothed curves and FPCA modes of variation

After recentering the Ripley curves relative to the empirical CSR reference, i.e. considering the profiles *L*_*i*_(*r*) − *L*_CSR_(*r*), where *L*_CSR_(*r*) denotes the mean curve obtained from random label permutations, and smoothing them using a B-spline basis representation (see Methods for details), the curves were analyzed using FPCA to characterize the main modes of variation between samples. The corresponding functional mean was calculated for descriptive purposes.

Beyond the mean, we aim to describe the main sources of variability between samples. We apply a functional principal component analysis (FPCA) to the smoothed curves of Fig 4 (a), which provides modes of variation around the mean. Fig 4 (b) shows the first two modes (denoted PC1(*r*) and PC2(*r*)). PC1 explains 98.9% of the variance, indicating that the variability between samples is essentially one-dimensional. This result is consistent with the observation that the recentered curves exhibit very similar shapes and differ mainly by their amplitude, particularly at larger distances (Fig 4 (a)). The second principal component PC2 explains a negligible proportion of variance (≈ 0.08%) and does not contribute meaningfully to the interpretation of spatial variability.

**Figure 4.**
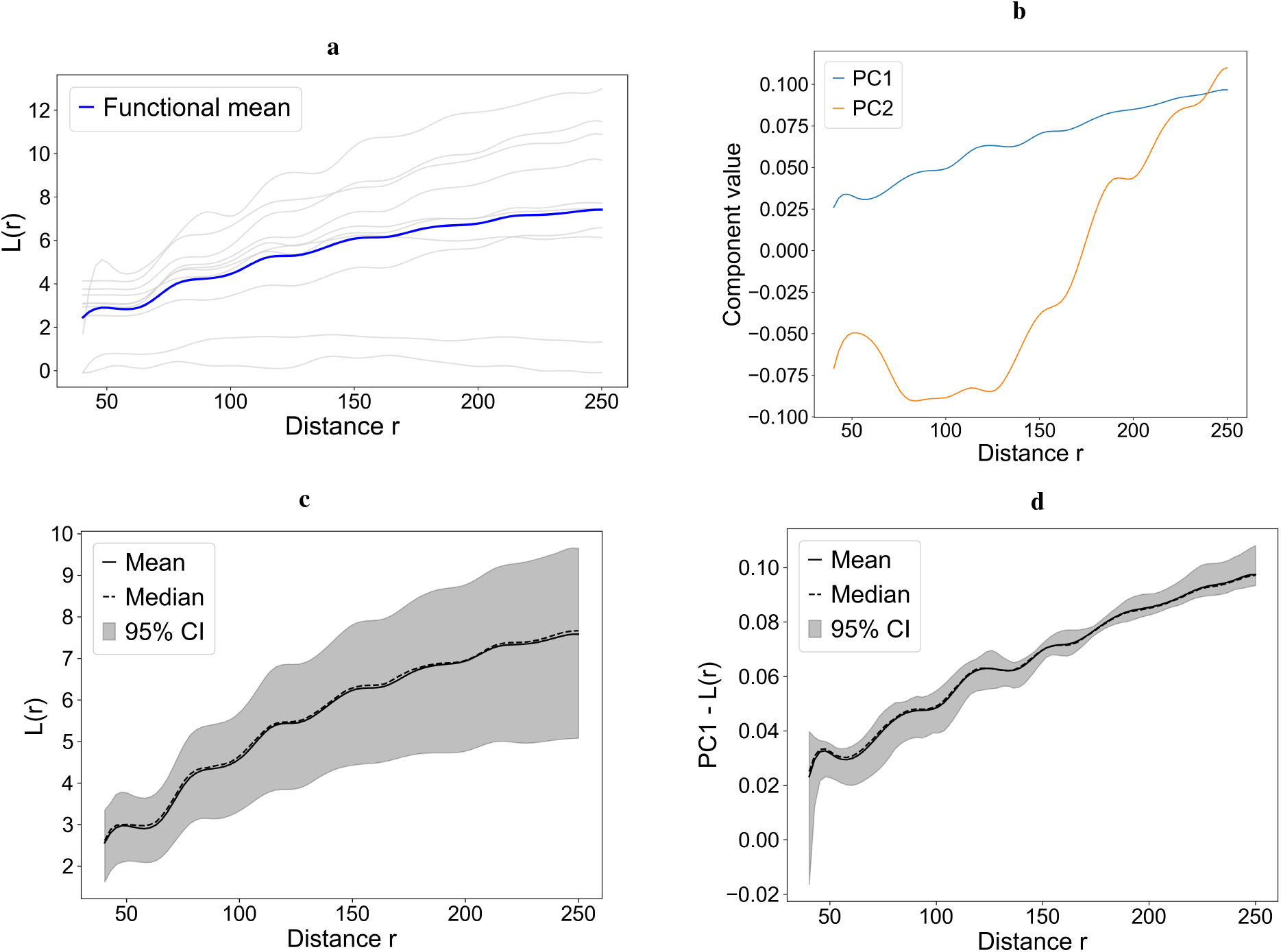
Smoothed recentered Ripley curves and functional modes of variation. **(a)** Smoothed and recentered Ripley *L*(*r*) curves for the samples (grey) together with the associated functional mean curve (blue). Curves were recentered relative to the empirical CSR reference described previously, restricted to distances *r* ≥ 40, and smoothed using a B-spline basis with 20 basis functions prior to FPCA. **(b)** First two functional principal components (FPCA) computed from the same recentered and smoothed Ripley *L*(*r*), illustrating the main shape variations of the *L*(*r*) curves around the mean. **(c)** Bootstrapped mean *L*(*r*) curve (mean, median and 95% CI over 100 resamples). **(d)** Stability of the first FPCA component (PC1) assessed by bootstrap (100 resamples), with mean, median and 95% CI.

In order to assess the stability of these results, we applied a bootstrap resampling procedure with replacement (100 repetitions). Fig 4 (c) shows the bootstrapped mean curve with its 95% confidence interval, whose width reflects the variability between samples, whereas Fig 4 (d) shows the stability of PC1 across resamples, with a relatively narrow 95% confidence interval given the small number of samples (n=10), suggesting that the dominant mode of variation identified by the FPCA is robust to resampling and not driven by a small subset of samples.

#### Functional projection and clustering of spatial aggregation profiles

To study inter-sample variability from a synthetic perspective, we now consider the space of FPCA scores. Each sample is represented by its coordinates on the plane defined by the first two functional principal components, which provide a lowdimensional representation of the recentered profiles introduced above. This representation is used to visualize the dispersion of spatial profiles between samples and to identify potential groups of similar behaviors through functional clustering.

The scores used in Fig 5 are obtained from a FPCA applied to these recentered curves. Prior to FPCA, the curves were smoothed and represented on a common domain.

**Figure 5.**
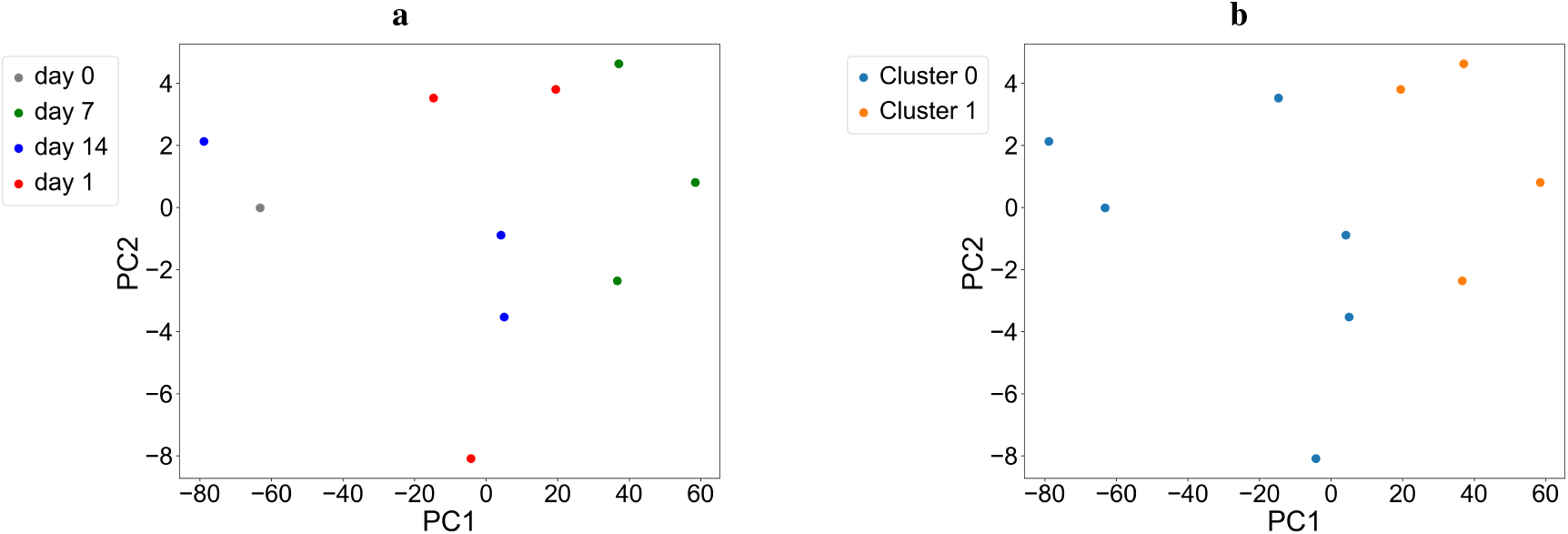
Functional analysis of Ripley *L*(*r*) curves using FPCA. **(a)** Projection of the samples onto the plane defined by the first two FPCA scores, computed from the smoothed recentered curves represented on a common B-spline basis over the interval *r* ∈ [40, 250]. Each point corresponds to one sample and is colored according to the day post-infarction. **(b)** Results of the functional clustering applied to the same curves, represented in the FPCA plane. Each color corresponds to a distinct functional cluster.

Each sample is then projected onto the plane defined by the first two principal components, which summarize the dominant modes of variation of the deviation from the CSR reference. The FPCA, and the clustering in the following, are performed on the interval *r* ∈ [40, 250], in accordance with the interpretation threshold retained for *L*(*r*), as discussed previously.

The distribution of samples in the FPCA plane in Fig 5 highlights differences between the spatial senescence profiles observed across samples. The dispersion of the points indicates that the Ripley curves do not all exhibit the same shape, reflecting variability in spatial aggregation profiles. However, the samples do not show a clear separation according to the time post-infarction: samples from different time points are intermingled, suggesting that temporal progression is not the primary driver of this variability within this dataset.

On the other hand, the representation of the clusters in the FPCA plane in Fig b reveals a clear partition between two functional groups resulting from the principal component decomposition. Cluster 0 is grouped around negative values of PC1, corresponding to curves exhibiting a shape characteristic of the first functional mode. Cluster 1 is distributed toward positive values of PC1 and more dispersed along PC2, reflecting greater variability.

#### Bootstrap validation of FPCA axes and scores

The classical analysis of an FPCA consists in interpreting the projection of the scores (PC1, PC2) as a reliable summary of inter-sample variability. However, when the number of curves is limited, recomputing the FPCA after a slight perturbation of the data may lead to a different basis, making the projections difficult to compare. In order to verify that the observed FPCA structure is not driven by the specific set of 10 curves available, we therefore performed a bootstrap with replacement by recomputing the FPCA at each iteration and evaluated the stability of the axes (cosine similarity of the components) as well as the variability of the associated FPCA scores.

As illustrated in Fig 6 (a), we assess this sensitivity by recomputing the FPCA on bootstrap sets of 10 curves sampled with replacement and aggregating all resulting projections. This interpretation is complemented by Fig 6 (B), which directly quantifies the stability of the axes through the cosine similarity of the components: PC1 appears almost invariant (mean/minimum similarity: 0.999/0.987) and PC2 remains globally stable on average (mean similarity: 0.900), although 19% of the bootstrap iterations exhibit a similarity below 0.9, reflecting occasional instability of this second mode. Within this stable functional reference frame, the point cloud obtained by bootstrap demonstrates that recomputing the FPCA after resampling does not significantly affect the projection: the scores remain globally organized in the same way. In other words, the method (axes and projection) is robust to small variations of the 10 curves, indicating that the observed trends do not depend on a particular sampling. After verifying by bootstrap the stability of the mean and median recentered curves curves in the distance space, we now aim to test the robustness of the same signal in the functional space subsequently used for FPCA. To do so, we fix in Fig 7 an FPCA learned from the 10 original samples. For each bootstrap resampling, we compute the corresponding mean profile and project it into this reference FPCA space, yielding the cloud of blue points. For comparison, the orange point represents the center of the original sample projections, obtained as the mean of the FPCA scores of the 10 samples. The dispersion of the bootstrap projections around this reference point is then used to assess the stability of the mean profile in the functional space.

**Figure 6.**
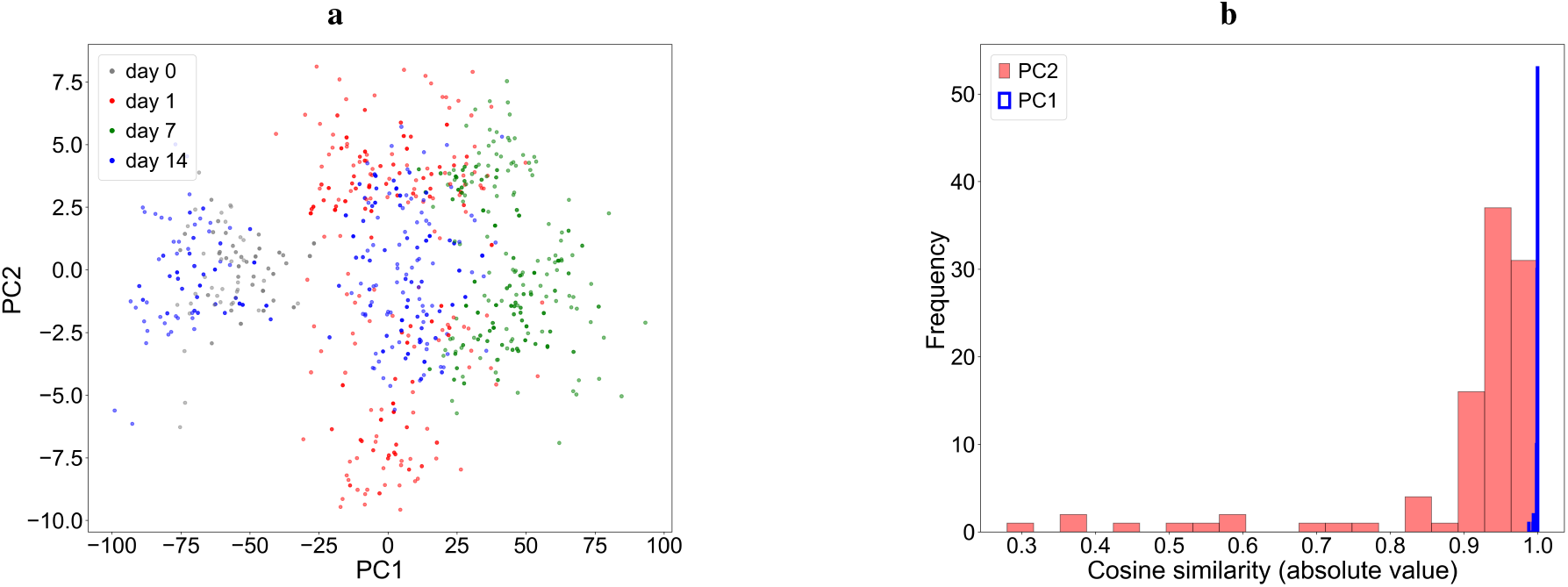
Stability of the FPCA projection under resampling (bootstrap). **(a)** Complete cloud of FPCA scores (PC1, PC2) obtained by recomputing the full FPCA on *B* = 100 bootstrap samples drawn with replacement and aggregating the resulting projections. **(b)** Distribution of cosine similarities (absolute value) between the FPCA components obtained for each bootstrap set and the reference components.

**Figure 7.**
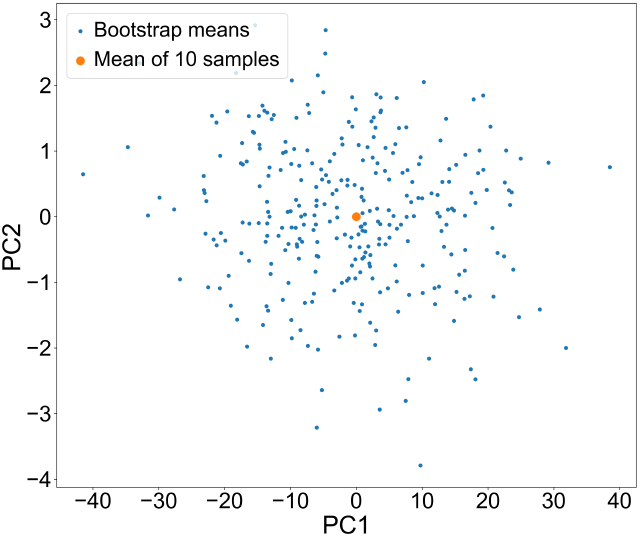
Bootstrap projection of mean recentered Ripley profiles. Bootstrap mean profiles (blue) projected onto the reference FPCA plane, compared with the center of the original FPCA scores (orange).

The fact that the cloud of bootstrap means remains grouped around the central point indicates that the mean profile does not shift strongly when the sampling is slightly modified: the mean estimator is therefore stable in the FPCA space. The dispersion observed around the center nevertheless reflects the heterogeneity of the spatial profiles across samples, consistent with the wide confidence bands obtained previously, but without any qualitative change in the overall trend.

### Beyond Ripley functions: pair-correlation functions

#### Pair-correlation functions and numerical estimation

Fig 8 shows the PCF *g*_sen,sen_(*r*) estimated for three individual samples corresponding to days 1, 7 and 14 post-infarction, selected to illustrate potential differences in spatial organization between post-infarction time points. As for the Ripley *L* function, the theoretical reference *g*(*r*) = 1 is not adapted to the discrete geometry of the Visium grid, and it is therefore replaced by an empirical reference obtained by permutation of the labels, following the same approach as previously.

**Figure 8.**
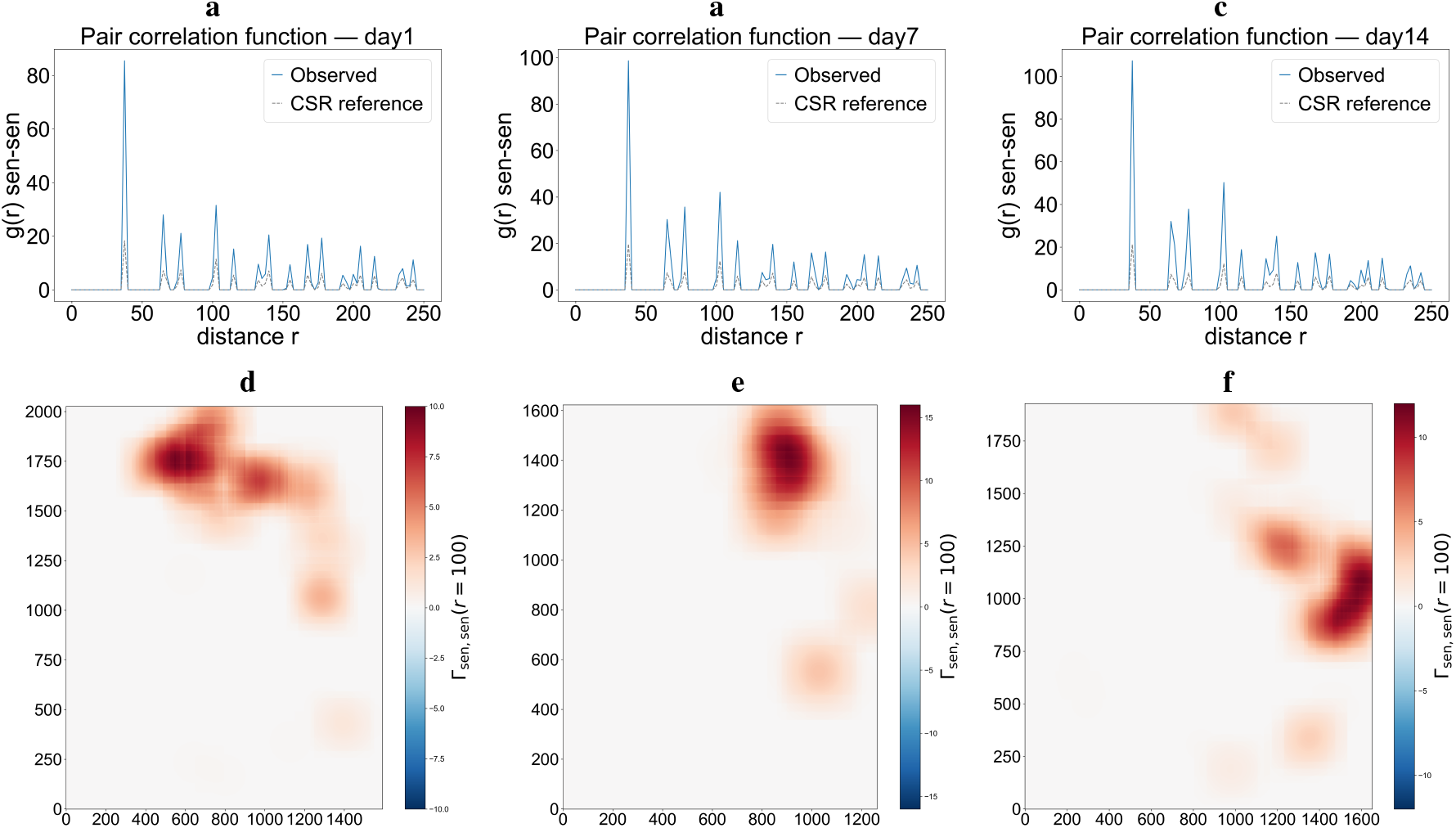
PCF of senescent spots and topographic correlation maps according to the day post-infarction. (**a-b-c**) Pair-correlation functions *g*(*r*) between senescent spots (sen-sen) for days 1, 7 and 14 post-infarction. The PCF was evaluated up to *r*_max_ = 250 pixels using annuli of width 2.5 pixels and a radial step of 2.5 pixels. The solid line represents the observed PCF and the dashed line the conditional CSR reference obtained from *B* = 199 random permutations of senescence labels. A detailed definition and interpretation of the PCF are provided in the Methods section. (**d-e-f**) Corresponding topographic correlation maps Γ_sen,sen_(*r* = 100) highlighting the spatial regions contributing to the senescent-senescent interaction at a distance of 100 pixels. Maps were computed using a Gaussian kernel (radius = 150 pixels, *σ* = 100 pixels) and a maximum correlation threshold of 5. A detailed description of the TCM construction is provided in the Methods section.

Sections from different days post-infarction exhibit a similar profile, dominated by a pronounced peak at short distances, suggesting a local spatial structure of senescent spots that remains stable over time. Indeed, the PCF shows a marked peak at short distance (*r* ≤ 50), with values of *g*(*r*) higher than the reference curve, indicating strong local aggregation of senescent spots.

Secondary peaks are observed at intermediate distances (*r* ≈ 70-150), suggesting a spatial organization structured at multiple scales. At larger distances (*r* ≥ 200), *g*(*r*) converges toward the reference curve, indicating that the aggregation of senescent spots is no longer distinguishable from a random distribution on the grid, as interactions between spots fade with distance. Moreover, the correlation maps reveal regions where the local density of senescent spots is 10 to 15 times higher than what would be expected under a purely random process, reflecting the presence of senescent aggregation hotspots within the tissue.

#### Functional analysis of pair-correlation functions

The PCFs *g*(*r*) are analyzed using FPCA in order to summarize the inter-sample variability of spatial interaction profiles and to identify their main directions of variation. Each PCF is represented using a B-spline basis with 17 basis functions in order to obtain a smooth functional representation over the distance domain [0, 250] and to preserve the main oscillatory patterns of the interaction profile.

The *g*(*r*) curves reveal aggregation patterns of senescent spots concentrated at short range, with an intensity that varies across samples. Fig 9 highlights that the first principal component, PC1 (70.6% of the variance), mainly reflects differences in PCF values between samples, with the largest differences observed at short distances. This suggests that most of the variability between samples is related to differences in the strength of short-range spatial interactions between senescent spots. The PC2 (4.6%) describes a secondary variation of the PCF profile, reflecting modulations of the main pattern already captured by the first component. Indeed, the proximity of the shapes associated with PC1 and PC2 indicates that the variability of the PCF is dominated by a common pattern, the second component reflecting adjustments of smaller amplitude around this pattern.

**Figure 9.**
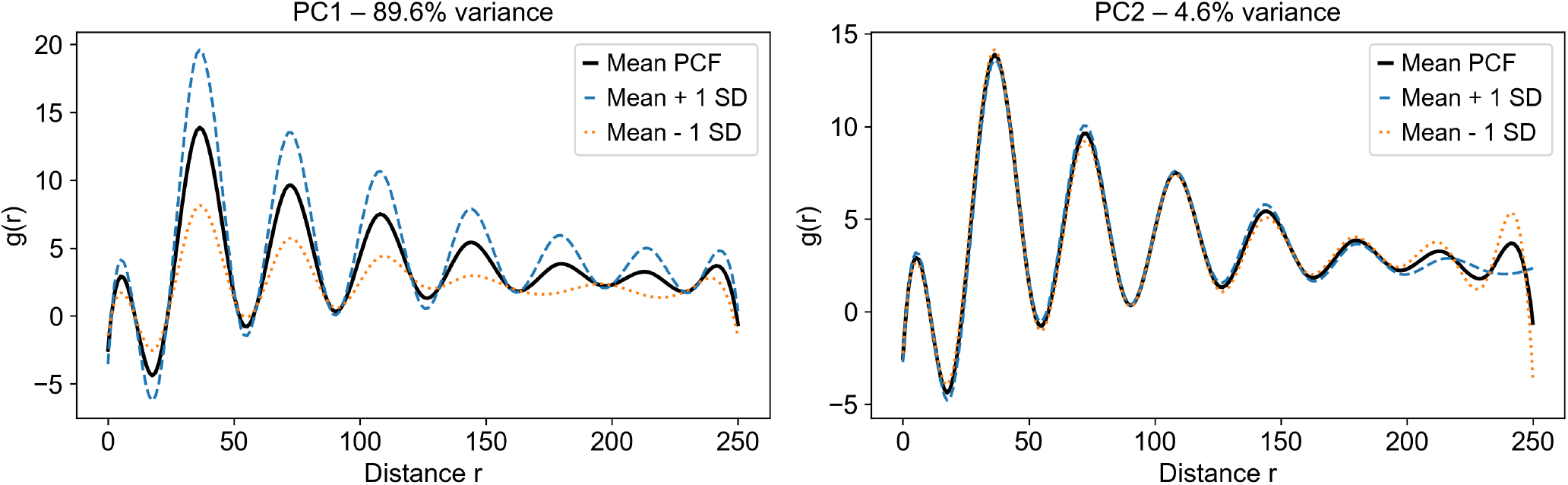
FPCA of pair-correlation functions between senescent spots. Mean pair-correlation function *g*(*r*) (solid line) together with the first (left) and second (right) functional principal modes of variation. Dashed lines represent the deformation of the mean function obtained by moving one standard deviation in the positive and negative directions of each component, thereby highlighting the dominant sources of variation.

## Discussion and Conclusion

A multi-scale analysis framework combining point process statistics (Ripley’s function *L*(*r*), PCF and TCM) with functional statistical approaches was used to characterize the spatial organization of senescent spots in Visium data. The application of this framework to mouse heart sections, in a context of myocardial infarction and across different time points, shows that senescence cannot be reduced to its abundance alone: the way senescent spots are structured within the tissue space exhibits variability, although the aggregation of these spots remains predominant. Indeed, by combining a measure of abundance (senescent burden) with a description of spatial structuring at different distances, our approach highlights regimes of aggregation or dispersion that would remain invisible using global indicators alone. More generally, this framework is directly transferable to other transcriptomic signatures and to other spatially resolved tissues, and provides a tool to relate molecular programs to their spatial contextualization within tissue.

In our Visium context, the comparison between a strictly theoretical CSR reference (homogeneous process, *L*(*r*) = *r*) and a reference estimated through permutations of labels highlights that the theoretical baseline may be inappropriate when the observed domain is constrained by the tissue mask, grid discretization, and the potential presence of excluded spots. In such situations, an apparent deviation from *L*(*r*) = *r* may reflect the effective geometry of the section rather than a biological interaction, which motivates the use of a conditional CSR reference obtained through permutation of the labels, preserving the admissible positions and the number of senescent spots.

However, this study remains limited by the restricted number of available samples. This constraint reduces the statistical power of inter-sample comparisons and prevents formally concluding the existence of systematic differences in spatial structuring between groups. In this context, the functional analyses and bootstrap procedures mainly aim to characterize inter-sample variability and the robustness of the observed trends, rather than to establish strict differential effects.

Moreover, the analysis is intrinsically constrained by the spatial resolution of the Visium technology. The arrangement of spots on a regular grid and their finite size impose a minimal distance threshold for interpretation, particularly visible around *r* ≈ 40 pixels. Below this threshold, spatial statistics mainly reflect the discretization of the support rather than biological interactions. Although this aspect has been explicitly considered in the interpretation of the results, it highlights the need for caution in multi-scale analysis and limits the ability to investigate very short-range interactions.

Beyond the methodological aspects, our results provide insight into the biological organization of senescence-associated transcriptional programs within damaged tissues. Across the analyzed heart sections, senescence-associated signals were consistently found to exhibit local aggregation rather than a random spatial distribution. This observation suggests that senescence-related programs do not emerge uniformly throughout the myocardium but instead concentrate within spatially restricted tissue regions. Such a pattern is consistent with the highly heterogeneous nature of post-infarction remodeling, where inflammatory responses, extracellular matrix remodeling, cell stress, and tissue damage are unevenly distributed across the affected tissue. Although the present study does not directly identify the mechanisms underlying these spatial patterns, the observed clustering supports the idea that senescence-associated programs are shaped by local tissue context. In this perspective, the separation observed by functional clustering in the FPCA in Fig 5 space may suggest the existence of distinct spatial organization profiles of senescence-associated programs, although this interpretation remains exploratory and cannot be directly assigned to specific biological regimes with the present dataset. More broadly, these findings suggest that heterogeneity in senescence should not be considered solely from a molecular perspective. In addition to the diversity of senescence-associated transcriptional states described in previous studies, our results indicate that senescence may also display a spatial dimension of heterogeneity within tissues. Characterizing how senescence-associated programs are organized in space may therefore contribute to a better understanding of their role in tissue remodeling, disease progression, and therapeutic response.

In conclusion, the multi-scale analysis of spatial organization using Ripley’s function *L*(*r*) highlights a robust tendency toward aggregation of senescent spots, observed in almost all of our 10 samples. This spatial structuring suggests that senescence-associated transcriptional programs, as inferred from SenePy-derived scores, do not emerge diffusely throughout the tissue but instead tend to concentrate within spatially restricted regions. Such aggregation patterns are compatible with the heterogeneous nature of post-infarction tissue remodeling and support the notion that the local tissue context contributes to the spatial organization of senescence-associated responses.

This conclusion is further supported by the analysis of pair-correlation functions, confirming the structured and multi-scale nature of senescent aggregation. Compared with a global indicator such as Moran’s index, which summarizes spatial autocorrelation into a single value, the combination of Ripley’s function and the PCF provides essential information about scale dependence by identifying the distances at which aggregation is most pronounced and by distinguishing potentially different spatial regimes depending on *r*.

This approach therefore provides a more refined and interpretable description of the spatial contextualization of senescence, and constitutes a solid basis for quantitatively comparing spatial profiles across post-infarction time points, while incorporating a null reference adapted to the real geometry of Visium sections.

The analyses presented in this work open several avenues for future investigation. First, validation on larger cohorts will be required to confirm the robustness and generalizability of the spatial patterns identified here. Extending this framework to additional spatial transcriptomics datasets, including human tissues and other pathological contexts such as fibrosis, cancer, or age-related diseases, such as those reported by Garbarino et al. [23] in human colorectal liver metastases, would help determine whether local aggregation of senescence-associated programs represents a general feature of tissue pathology. Furthermore, integrating spatial organization with molecular, cellular, or microenvironmental information could provide deeper insight into the biological mechanisms underlying the emergence and maintenance of senescence-associated regions within tissues.

On the other hand, the Ripley functions used here rely on a uniform weighting of neighbors, which constitutes a simplification: a distant point contributes as much as a nearby point. A natural extension would consist in adopting adaptive kernels, where nearby neighbors are assigned a greater weight than distant neighbors, making the analysis more faithful to the biological reality of local interactions. This approach, developed in particular by Hohl et al. [24], would allow the construction of local versions of the Ripley function better suited to the spatial heterogeneity of tissues.

## Acknowledgements

We thank Alexandre Poulain for his availability, his constructive feedback, and his careful reviews, as well for his expertise in mathematics, which greatly contributed to strengthening the quality and rigor of this work.

We thank Ko T, Hatsuse S, Nomura S, Aburatani H, Komuro I (GSE176092, NCBI GEO) for making the public data used in this study available.

We thank the open-source community for the development and maintenance of the tools used (Scanpy/AnnData, Squidpy, FDApy, SciPy).

The authors gratefully acknowledge the CNRS for funding C.V.’s PhD fellowship and also thank the Inria DATAVERS project-team and ONCOLille for their scientific support and for providing a quality research environment.

All figures and illustrations presented in this work were created by the authors.

## Author contributions statement

C. V. performed the analyses, developed the computational framework, generated the figures, and led the writing of the manuscript. S.D. supervised the research, contributed to the interpretation of the results and contributed to manuscript writing and revision. V.D. provided biological expertise, contributed to the interpretation of the findings and contributed to manuscript writing and revision. All authors approved the final manuscript.

## Supplementary information

Supplementary Tables S1-S2 accompany this paper.

## Data Availability

All figures were created by the authors. Figure 1 was designed using BioRender (https://www.biorender.com/). Figure 2.a includes representative images from the publicly available dataset GSE176092 described by Yamada et al. [18]. All other figures were generated from analyses performed in Python (version 3.12, https://www.python.org/). The datasets analysed during the current study are publicly available in the NCBI Gene Expression Omnibus (GEO) repository under accession number GSE176092 https://www.ncbi.nlm.nih.gov/geo/query/acc.cgi?acc=GSE176092 by Yamada et al. [18].

## References

1. Campisi, J. Aging, cellular senescence, and cancer. Annu. Rev. Physiol. 75, 685–705, 10.1146/annurev-physiol-030212-183653 (2013).

2. Childs, B. G., Durik, M., Baker, D. J. & van Deursen, J. M. Cellular senescence in aging and age-related disease: from mechanisms to therapy. Nat. Medicine 21, 1424–1435, 10.1038/nm.4000 (2015).

3. Gorgoulis, V. et al. Cellular senescence: Defining a path forward. Nat. Rev. Mol. Cell Biol. 20, 671–691, 10.1038/s41580-019-0169-z (2019).

4. Hernandez-Segura, A., Nehme, J. & Demaria, M. Hallmarks of cellular senescence. Trends Cell Biol. 28, 436–453, 10.1016/j.tcb.2018.02.001 (2018).

5. Prasanth, M. I., Brimson, J. M. & Tencomnao, T. Virus-induced senescence: a therapeutic target for mitigating severe progression of sars-cov-2. Expert. Rev. Mol. Medicine 25, 263–267, 10.1080/14728222.2023.2218033 (2023).

6. Sanborn, M. A., Wang, X., Gao, S., Dai, Y. & Rehman, J. Unveiling the cell-type-specific landscape of cellular senescence through single-cell transcriptomics using senepy. Nat. Commun. 16, 10.1038/s41467-025-57047-7 (2025).

7. Loison, I. et al. O-glcnacylation inhibition redirects the response of colon cancer cells to chemotherapy from senescence to apoptosis. Cell Death Dis. 15, 10.1038/s41419-024-07131-5 (2024).

8. Xu, M. et al. Senolytics improve physical function and increase lifespan in old age. Nat. Medicine 24, 1246–1256, 10.1038/s41591-018-0092-9 (2018).

9. Zhu, Y. et al. The achilles’ heel of senescent cells: from transcriptome to senolytic drugs. Aging Cell 14, 644–658, 10.1111/acel.12344 (2015).

10. Wang, B., Kohli, J. & Demaria, M. Senescent cells in cancer therapy: Friends or foes? Trends Cancer 6, 838–857, 10.1016/j.trecan.2020.05.004 (2020).

11. Ripley, B. D. The second-order analysis of stationary point processes. J. Appl. Probab. 13, 255–266, 10.2307/3212829 (1976).

12. Sampaio, W. B., Diniz, E. M., Silva, A. C. & de Paiva, A. C. Detection of masses in mammograms using cellular neural networks, hidden markov models and ripley’s k function. Proceedings of the 16th International Conference on Systems, Signals and Image Processing, 10.1109/IWSSIP.2009.5367756 (IEEE, 2009).

13. Wilson, C. et al. Tumor immune cell clustering and its association with survival in african american women with ovarian cancer. PLOS Comput. Biol. 18, e1009900, 10.1371/journal.pcbi.1009900 (2022).

14. Krishnan, S. N., Mohammed, S., Frankel, T. L. & Rao, A. Gawrdenmap: a quantitative framework to study the local variation in cell–cell interactions in pancreatic disease subtypes. Sci. Reports 12, 3708, 10.1038/s41598-022-06602-z (2022).

15. Ramsay, J. O. & Silverman, B. W. Functional Data Analysis. Springer Series in Statistics (Springer, 1997).

16. Wrobel, J. et al. mxfda: a comprehensive toolkit for functional data analysis of single-cell spatial data. Bioinforma. Adv. 4, vbae155, 10.1093/bioadv/vbae155 (2024).

17. Dabo-Niang, S. & Frévent, C. Uncovering data across continua: An introduction to functional data analysis. arXiv preprint arXiv:2404.16598 (2024).

18. Yamada, S. et al. Spatiotemporal transcriptome analysis reveals critical roles for mechano-sensing genes at the border zone in remodeling after myocardial infarction. Nat. Cardiovasc. Res. 1, 1072–1083, 10.1038/s44161-022-00165-w (2022).

19. Wolf, F. A., Angerer, P. & Theis, F. J. Scanpy: large-scale single-cell gene expression data analysis. Genome Biol. 19, 15, 10.1186/s13059-017-1382-0 (2018).

20. Moran, P. A. P. Notes on continuous stochastic phenomena. Biometrika 37, 17–23, 10.1093/biomet/37.1-2.17 (1950).

21. Marcon, E., Floch, J.-M. & Puech, F. Spatial distribution of points. In Loonis, V. & de Bellefon, M.-P. (eds.) Handbook of Spatial Analysis: Theory and Application with R, 71–112 (Insee and Eurostat, (2018)).

22. Bull, J. A., Mulholland, E. J., Leedham, S. J. & Byrne, H. M. Extended correlation functions for spatial analysis of multiplex imaging data. Biol. Imaging 4, e2, 10.1017/S2633903X24000011 (2024).

23. Garbarino, O. et al. Spatial resolution of cellular senescence dynamics in human colorectal liver metastasis. Aging Cell 22, e13853, 10.1111/acel.13853 (2023).

24. Hohl, A. & Chen, P. Spatiotemporal simulation: local ripley’s k function parameterizes adaptive kernel density estimation. In Proceedings of the 2nd ACM SIGSPATIAL International Workshop on Spatial Simulation, 10.1145/3356470.3365528 (2019).

