## Supplementary Information for "Multiscale Analysis of Cellular Senescence through Ripley’s Functions and Functional Statistics"

To complement the visual exploration of spatial organization presented in Figure 1 of the main manuscript, Supplementary Tables S1 and S2 summarize the statistical significance of the observed spatial structures using Ripley’s statistic and Moran’s index, respectively.

For Ripley’s statistic, exact p-values were obtained from a one-sided permutation test based on (B=199) random permutations of the senescence labels. The null hypothesis corresponds to a random spatial distribution of senescent spots, while the alternative hypothesis corresponds to a stronger spatial aggregation than expected under complete spatial randomness. For Moran’s index, p-values were obtained from a one-sided permutation test implemented in *PySAL* . Since both analyses rely on permutation procedures, they do not require any assumption of normality. Statistical significance was assessed at the α=0.05 level.

### Supplementary Table S1. Permutation p-values of the Ripley statistic computed at four distances for the 10 samples using B = 199 permutations.

| Data | Day | p(r=50) | p(r=100) | p(r=150) | p(r=200) |
| --- | --- | --- | --- | --- | --- |
| GSM5355663 | Sham | 0.040 | 0.005 | 0.005 | 0.020 |
| GSM5943189 | 1 | 0.005 | 0.005 | 0.005 | 0.005 |
| GSM5943190 | 1 | 0.005 | 0.005 | 0.005 | 0.005 |
| GSM5943191 | 1 | 0.005 | 0.005 | 0.005 | 0.005 |
| GSM5355666 | 7 | 0.005 | 0.005 | 0.005 | 0.005 |
| GSM5943192 | 7 | 0.005 | 0.005 | 0.005 | 0.005 |
| GSM5943193 | 7 | 0.005 | 0.005 | 0.005 | 0.005 |
| GSM5355668 | 14 | 0.595 | 0.445 | 0.165 | 0.345 |
| GSM5943194 | 14 | 0.005 | 0.005 | 0.005 | 0.005 |
| GSM5943195 | 14 | 0.005 | 0.005 | 0.005 | 0.005 |

### Supplementary Table S2. Moran's I index and associated permutation p-value for each sample.

| Data | Day | Moran's I | p-value |
| --- | --- | --- | --- |
| GSM5355663 | Sham | 0.0317 | 0.014 |
| GSM5943189 | 1 | 0.1566 | 0.001 |
| GSM5943190 | 1 | 0.1845 | 0.001 |
| GSM5943191 | 1 | 0.2103 | 0.001 |
| GSM5355666 | 7 | 0.3063 | 0.001 |
| GSM5943192 | 7 | 0.2641 | 0.001 |
| GSM5943193 | 7 | 0.2705 | 0.001 |
| GSM5355668 | 14 | -0.0063 | 0.430 |
| GSM5943194 | 14 | 0.2288 | 0.001 |
| GSM5943195 | 14 | 0.2647 | 0.001 |

Permutation p-values were computed using the empirical null distribution obtained from
B = 199 label permutations.

### Supplementary Figure S1. Histological images and spatial senescence score maps for all 10 tissue sections.

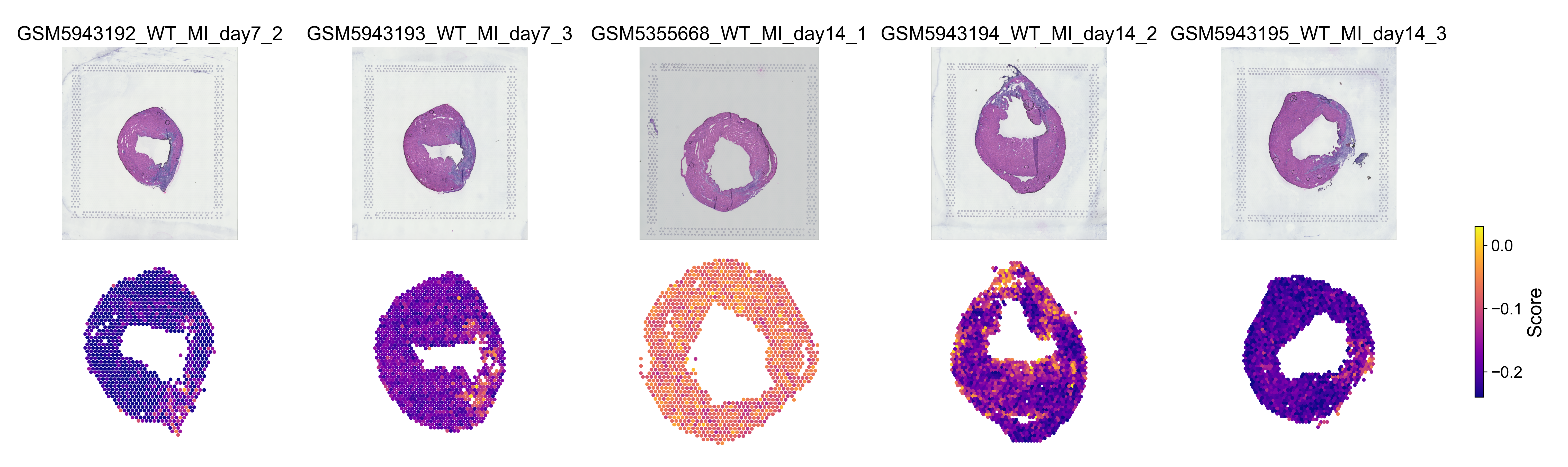

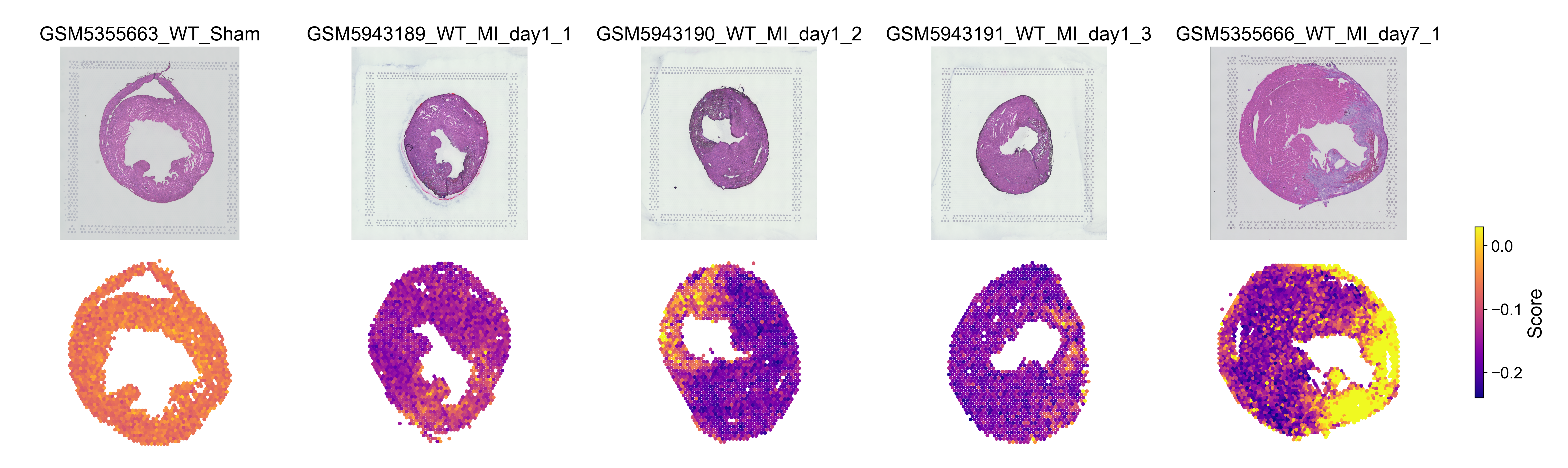

For each sample, the histological image of the tissue section is shown above the corresponding spatial senescence score map for the cardiac myocyte hub identified using SenePy. Samples are ordered chronologically: Sham, day 1, day 7, and day 14 post-infarction, with three replicates at each post-infarction time point. Senescence scores across Visium spots are displayed using a fixed plasma color scale shared across all samples, enabling direct visual comparison.
